# ClinPhen extracts and prioritizes patient phenotypes directly from medical records to accelerate genetic disease diagnosis

**DOI:** 10.1101/362111

**Authors:** Cole A. Deisseroth, Johannes Birgmeier, Ethan E. Bodle, Jonathan A. Bernstein, Gill Bejerano

## Abstract

**Purpose:** Severe genetic diseases affect 7 million births per year, worldwide. Diagnosing these diseases is necessary for optimal care, but it can involve the manual evaluation of hundreds of genetic variants per case, with many variants taking an hour to evaluate. Automatic gene-ranking approaches shorten this process by reporting which of the genes containing variants are most likely to be causing the patient’s symptoms. To use these tools, busy clinicians must manually encode patient phenotypes, which is a cumbersome and imprecise process. With 60 million patients expected to be sequenced in the next 7 years, a fast alternative to manual phenotype extraction from the clinical notes in patients’ medical records will become necessary.

**Methods:** We introduce ClinPhen: a fast, high-accuracy tool that automatically converts the clinical notes into a prioritized list of patient symptoms using HPO terms.

**Results:** ClinPhen shows superior accuracy to existing phenotype extractors, and when paired with a gene-ranking tool it significantly improve the latter’s performance.

**Conclusion:** Compared to manual phenotype extraction, ClinPhen saves more than 5 hours per case in Mendelian diagnosis alone. Summing over millions of forthcoming cases whose medical notes await phenotype encoding, ClinPhen makes a substantial contribution towards ending all patients’ diagnostic odyssey.

## Introduction

Every year, 7 million children worldwide^1^ are born with rare genetic diseases. Diagnosing these conditions currently involves determining which (if any) of numerous genetic variants (100-300 coding variants alone, if there are no sequenced relatives) is causing the patient’s symptoms. This is done by spending an average of 54 minutes evaluating each variant^2^ until the causative one is identified (Figure 1). As sequencing technology becomes more time- and cost-efficient, the number of clinical applications skyrockets, to the point that over 60 million patients are expected to be sequenced by 2025^3^. As more patients with genetic diseases are sequenced, the procedure of manually evaluating the possible causative variants for each patient creates a bottleneck in the diagnostic process.

**Figure 1.**
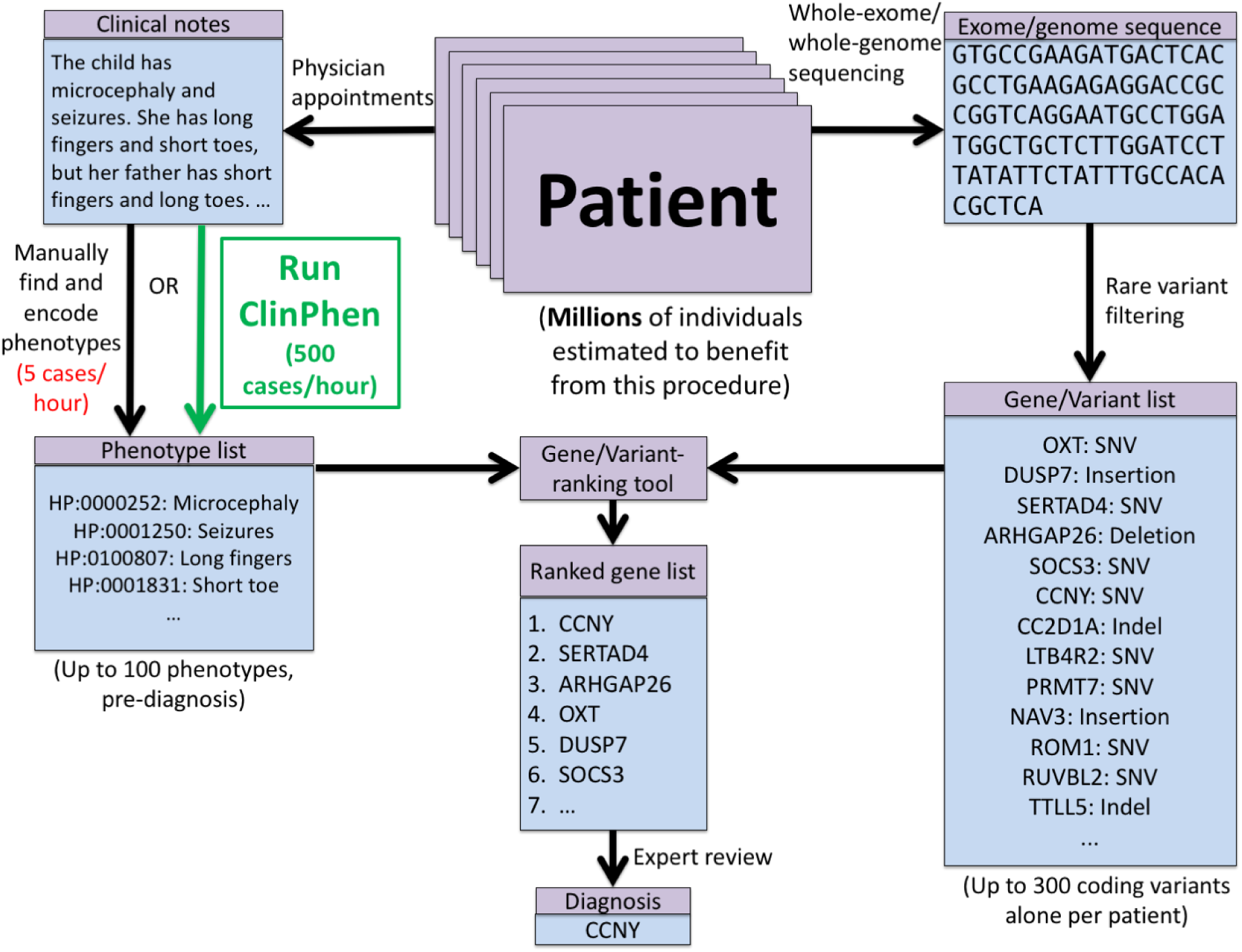
Steps to diagnose a patient with a Mendelian disease using automated gene-ranking algorithms. The patient’s genotypic information is encoded using standard formats (Variant Call Format (VCF) file, candidate causative gene list) and a list of patient phenotypes encoded as ontology terms. Extensive tool support exists for obtaining candidate causative variants and genes from an exome sequence. Tool support for obtaining an appropriate list of encoded patient phenotypes from the patient’s clinical notes is limited. Encoded phenotypes are currently acquired by manually reading through the patient’s clinical notes and recording the phenotypes found as their IDs in a phenotype ontology. We introduce ClinPhen, a tool that automates phenotype extraction from clinical notes, optimized for accelerated patient diagnosis.

Although a clinician must make the final decision on the diagnosis, the process leading up to it can be greatly expedited by computational tools. Phrank^4^, hiPhive^5^, Phive^6^, PhenIX^7^, and other automatic gene-ranking tools^8–16^ aim to speed up the process of evaluating potentially causative genes. These algorithms require as input a list of symptoms encoded as terms from an existing phenotype ontology (notably, the Human Phenotype Ontology, or HPO^17^). In addition, they require a list of genes containing potentially deleterious genetic variants. These tools automatically rank the list of candidate genes using their own estimates of each gene’s likelihood of causing the patient’s disease. Consequently, the clinician can reach a diagnosis faster by going down the ranked list, evaluating the candidate genes until the causative gene is identified.

The process of annotating a patient’s genetic variants and filtering them to a list of candidate genes is greatly facilitated by tools such as ANNOVAR^18^, VAAST^19^, VEP^20^, and snpEff^21^. However, comparable tools for automatically encoding phenotypes mentioned in the patient’s clinical notes into phenotype ontology terms are lacking. While gene-ranking tools can considerably shorten the lengthy manual review of a gene list to reach a diagnosis^2^, their ability to do so depends on the phenotypes used (see below). The manual encoding of phenotypes is a slow and unstructured process, making gene-ranking tools difficult for clinicians to adopt.

Existing Natural Language Processing tools that identify patient phenotypes were not designed to accelerate the diagnosis of Mendelian diseases^22–28^. Many of these tools only look for indications of specific subsets of phenotypes or diseases^27,29,30^. Others report and encode all of the phenotypes they can find—including negated phenotypes (“The patient does *not* have symptom X”), findings in family members (“The patient’s *mother* has symptom X”), and phenotypes mentioned in the discussion of a differential diagnosis (“Patients with disease W often have symptoms X, Y, and Z”)—requiring manual review of the clinical notes to remove the out-of-context phenotypes^24,28^. Two general-purpose phenotype extractors, cTAKES^26^ and MetaMap^22^, aim to extract only the phenotypes that apply to the patient, thus returning a list of patient phenotypes that are ready to run through an automatic gene-ranking tool.

cTAKES and MetaMap, however, have not been optimized for a clinical genetics workflow. They have relatively slow runtimes and suboptimal accuracy (see below). Importantly, they do not indicate which phenotypes may be more useful in establishing a diagnosis. A patient’s clinical notes taken before the diagnosis can have over 100 phenotypes mentioned about the patient, but clinicians do not usually list all of them when trying to diagnose. Instead, they list only the ones that they think will be the most useful in diagnosing the patient.

Here, we introduce ClinPhen: a fast, easy-to-use, high-precision, and high-sensitivity alternative to existing phenotype extractors. ClinPhen scans through a patient’s clinical notes in seconds, and returns phenotypes that help gene-ranking tools rank the causative gene higher than they would with manually-identified phenotypes. Using a cohort of diagnosed patients, we show how to accelerate the diagnosis of Mendelian diseases by letting gene-ranking tools run directly on phenotypes extracted from the clinical notes by ClinPhen.

## Materials and Methods

### Overview of ClinPhen

ClinPhen is an algorithm for fast, accurate phenotype extraction from clinical notes (Figure 2). ClinPhen extracts phenotypes encountered in the free-text notes and translates them into terms from the Human Phenotype Ontology (HPO), a structured database containing 29,107 names and synonyms of 12,486 human disease phenotypes. ClinPhen uses the UMLS Metathesaurus^31^ and the Monarch Initiative^32^ to expand HPO’s synonym list from 29,107 to 69,690 synonyms for the same 12,486 phenotypes.

**Figure 2.**
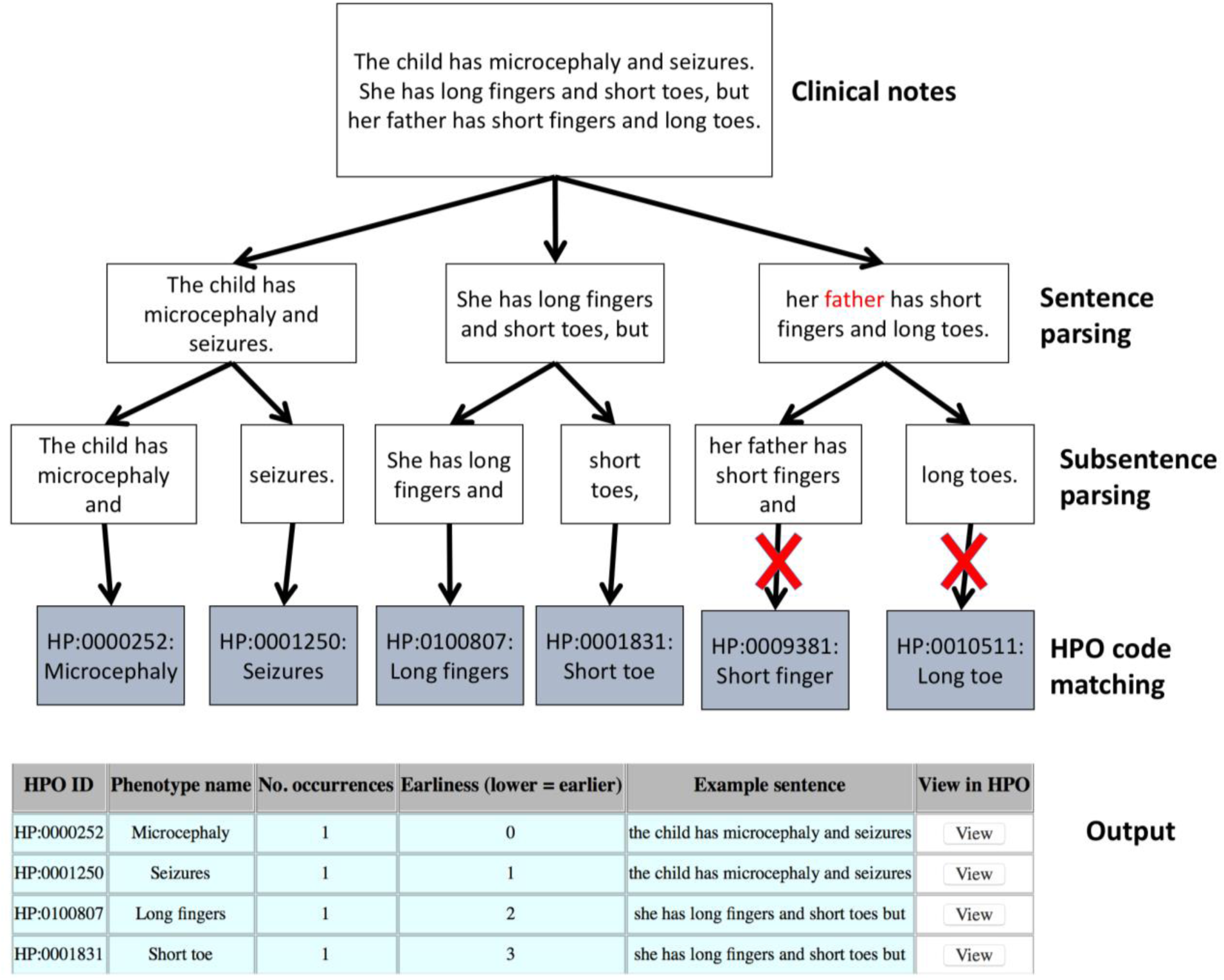
ClinPhen sentence analysis process. ClinPhen splits the clinical notes into sentences, and those sentences into subsentences. It then finds phenotypes whose synonyms appear in the subsentences. A high-precision, high-sensitivity, rule-based natural-language-processing system decides which phenotypes correspond to true mentions and which are false positives. Since the third sentence contains the flag word “father”, for instance, it is assumed that this sentence does not refer to the patient, and any phenotype synonyms found in the sentence will not be considered. ClinPhen sorts the identified phenotypes first by how many times they appeared in the notes (descending), then by the index of the first subsentence in which they were found (ascending), and then by HPO ID (ascending).

To extract HPO terms from the clinical notes, ClinPhen proceeds as follows: first, the free text is broken into sentences, subsentences, and words. ClinPhen “lemmatizes” the words using the Natural Language Toolkit (NLTK) Lemmatizer^33^. “Lemmatization” refers to a process in which inflected forms of words are replaced with normalized forms. The word “lesions”, for instance, is lemmatized to “lesion”. This allows ClinPhen to accept phrases such as “localized skin lesions” (plural) to match the name “localized skin lesion” (singular).

Subsequently, ClinPhen uses a custom rule-based heuristic to match subsentences against phenotype names and synonyms (Online Methods). ClinPhen does not look for continuous phrases, but instead checks if the subsentence contains all words in the given synonym. As an example, “Hands are large” will be a valid mention of the phenotype “Large hands”. For efficiency, ClinPhen does not search the entire collection of notes for each phenotype. Instead, it uses a hash table^34^ to pass over the notes once, record which words appeared in which subsentences, and then be able to quickly identify which subsentences in the notes contain all of the words in a given phenotype name.

After identification of phenotype terms in free text, ClinPhen uses a rule-based Natural-Language-Processing (NLP) framework to decide how a phenotype mention should be interpreted. The framework relies on an extensive list of keywords to decide if a mentioned phenotype applies to the patient. If, for instance, a sentence contains words such as “not” or “mother”, then ClinPhen assumes any phenotypes mentioned in the sentence do not apply to the patient.

For each HPO phenotype, ClinPhen counts the number of valid occurrences in the clinical notes, and saves where in the notes it first appears. ClinPhen returns a sorted list of all HPO phenotypes found, with the most- and earliest-mentioned phenotypes at the top (Figure 2).

### ClinPhen extracts the most accurate phenotype sets

#### Real patient cases used to improve and test ClinPhen

ClinPhen was trained and tested on patient data obtained from the clinical genetics service at Stanford Children’s Health (SCH). Two sets of patient data were used. The Training set consisted of the clinical notes of 20 patients with presumed but undiagnosed genetic diseases, and was used to improve the accuracy of ClinPhen. The Test set consisted of the clinical notes, genetic data, and diagnoses of all available (24) patients who had clinical notes from genetics
and/or pediatrics, were diagnosed with a genetic disease by Medical Genetics at the Lucille Packard Children’s Hospital (LPCH) in Stanford, and consented for research use. This set was used to test the accuracy of ClinPhen, as well as the performance of gene-ranking tools when using ClinPhen’s phenotypes. For all experiments, only notes created by the clinical genetics and pediatrics services at LPCH before the patient’s diagnosis were used.

#### Ontology and thesauruses

ClinPhen uses the Directed Acyclic Graph (DAG) of phenotypic abnormalities provided by the Human Phenotype Ontology (HPO)^17^ The HPO DAG is a large collection of phenotypes, where the more-general “parent” phenotypes are linked to their more-specific subcategories, or “child” phenotypes. “Generalized tonic-clonic seizures”, for instance, is a child of “Generalized seizures”, which is a child of “Seizures”. HPO also has a list of synonyms for every phenotype. “Seizures”, “Seizure”, and “Epilepsy”, for instance, all correspond to the same phenotype, represented by the ID HP:0001250. ClinPhen looks for these synonyms in the clinical notes to determine if the phenotype is mentioned. All phenotypes descending from the node “Phenotypic Abnormality” (HP:000018) are considered. Since many of the widely-used synonyms for phenotypes are not yet included in HPO, ClinPhen supplements HPO’s thesaurus using the metathesauruses provided by the Monarch Initiative^32^ and the Unified Medical Language System (UMLS)^31^. These two databases provide additional synonyms for the 12,486 HPO phenotypes (from the July 2017 release), and expand ClinPhen’s vocabulary from 29,107 to 69,690 phenotype synonyms.

#### Sentence and subsentence splitting and flagging

To extract phenotypes from clinical notes, ClinPhen splits the notes into sentences using a set of sentence delimiters. Each sentence is split into a list of subsentences using a set of subsentence delimiters. ClinPhen additionally records each sentence’s “flags”: words that indicate that a phenotype mention may not apply to the patient. For phenotypes such as “Negative chorea”, ClinPhen will count the phenotype as validly mentioned, even though the sentence contains the flag word “negative”.

#### Training flags and delimiters used by ClinPhen

We used the Training set to manually determine which words and characters are best used as flags or delimiters. Phenotypes from the clinical notes of these 20 patients were extracted, once manually, and once by ClinPhen. The flags and delimiters used by ClinPhen were optimized so that ClinPhen’s phenotypes would be as similar as possible to those found manually.

The set of sentence delimiters after training consisted of periods, bullet points, tabs, semicolons, newlines (Only during a second pass, in which ClinPhen checks for phenotype lists in the format “criterion: value”, which are common in EMRs, and often lack periods or other sentence delimiters), and the words “but”, “except”, “however”, and “though”. The set of subsentence delimiters after training consisted of commas, colons, and the word “and”. The set of flags included words that indicate that the mentioned phenotype applies to a family member, not the patient (cousin, parent, mom, mother, dad, father, grandmother, grandfather, grandparent, family, brother, sister, sibling, uncle, aunt, nephew, niece, son, daughter, grandchild); words that directly negate the mentioned phenotype (no, not, none, negative, non, never, normal); and words that indicate that the phenotypes are mentioned as part of a differential diagnosis (associated, gene, recessive, dominant, variant, cause, patients, literature, individuals).

#### Accuracy Testing

To test the accuracy of the extracted phenotypes, we compared the *All* set to the phenotypes returned by ClinPhen across all of the Test Patients. Due to the nature of the HPO DAG, the presence of a phenotype in a patient implies the presence of all ancestor phenotypes in the patient. For instance, the term “Seizures” is an ancestor node of the term “Grand mal seizures”: if a patient presents with grand mal seizures, then the patient also presents with seizures. The “closure” of a set of HPO terms *S* consists of *S* plus all ancestors of the terms in *S* up to “Phenotypic Abnormality” (HP:0000118). We compared the extracted phenotypes to the true phenotypes using the closures of the two sets.

For each of the Test Patients, we found the closure of the *All* set and that of the phenotype set returned by ClinPhen. True positives were defined as the nodes that are present in both the *All* and ClinPhen closures. False positives were defined as the nodes only present in the ClinPhen closure. False negatives were defined as the nodes only present in the *All* closure. The standard definitions of precision and sensitivity were used, as given by equations (1) and (2).

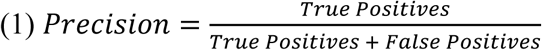

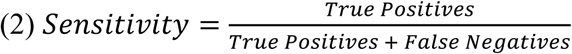

We calculated the 95% confidence interval around the average precision using equation (3).

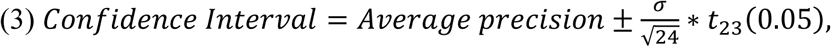

where σ is the standard deviation of the precision across all 24 Test patients and t_23_ is the inverse of Student’s t distribution with 23 degrees of freedom. The confidence interval around the average sensitivity was calculated similarly.

Since the phenotype extractors cTAKES and MetaMap output UMLS terms, not HPO terms, we translated the output of these tools to HPO using the UMLS Metathesaurus, which matches UMLS phenotypes to synonymous HPO phenotypes.

### Automatic extraction of phenotypes accelerates Mendelian disease diagnosis

Per conventions in the field ^35^ we used Exome Aggregation Consortium^36^ and 1000 Genomes Project^37^ frequency data to produce a fixed list of candidate causative genes per patient. The lists were used to compare all gene-ranking methods. Please see Supplementary Methods and Materials for the details.

### ClinPhen consistently runs in less than 10 seconds

As mentioned above, for each of the 24 Test patients, a licensed physician timed himself reading through the clinical notes, manually extracting the phenotypes that he considered useful for diagnosis and finding their matching HPO terms, thus creating the *Clinician* set. The times taken by the physician served as reference points for how long a clinician would take to manually extract phenotypes from clinical notes. We also timed each of the automatic phenotype extractors when running them on the same clinical notes.

## Results

### ClinPhen extracts the most accurate phenotype sets

A phenotype extractor can best help with disease diagnosis if its extracted phenotypes accurately reflect the patient’s symptoms. We compare the accuracy of 3 tools that can automatically extract phenotypes from clinical notes: ClinPhen, cTAKES, and MetaMap.

To determine which extractor is the most accurate, we used a Test set of 24 real patients diagnosed with Mendelian diseases. Each patient was associated with next-generation sequencing data, a diagnosis (including a single causative gene), and the clinical notes created before the diagnosis. For each Test patient, we produced a gold-standard phenotype set called the *All* set: A licensed physician and a non-physician independently extracted phenotypes from the clinical notes. The physician recorded only the phenotypes that he considered useful for diagnosis (those that are more likely to pertain to a genetic disease, such as skeletal abnormalities, as opposed to allergies) to generate the *Clinician* phenotype set. The non-physician recorded all of the phenotypes he found, regardless of predicted usefulness. The physician then verified the non-physician’s identified phenotypes to be correctly interpreted and applicable to the patient. These verified phenotypes, plus those in the *Clinician* set, made up the *All* phenotype set. We ran each automatic phenotype extractor on the patient’s clinical notes, and measured the extractor’s precision and sensitivity by comparing the extracted phenotypes to the *All* set.

Across the 24 Test patients, cTAKES had an average precision of 57%, and MetaMap had an average precision of 56%. ClinPhen had a precision of 70%, significantly higher than that of cTAKES or MetaMap (both p-values < 3.0^∗^10^-7^; Wilcoxon signed rank text).

cTAKES had an average phenotype sensitivity of 57%, and MetaMap had an average phenotype sensitivity of 71%. ClinPhen had an average phenotype sensitivity of 72%, significantly higher than that of cTAKES (p<3.0^∗^10^-7^), and slightly higher than that of MetaMap (Figure 3a).

**Figure 3.**
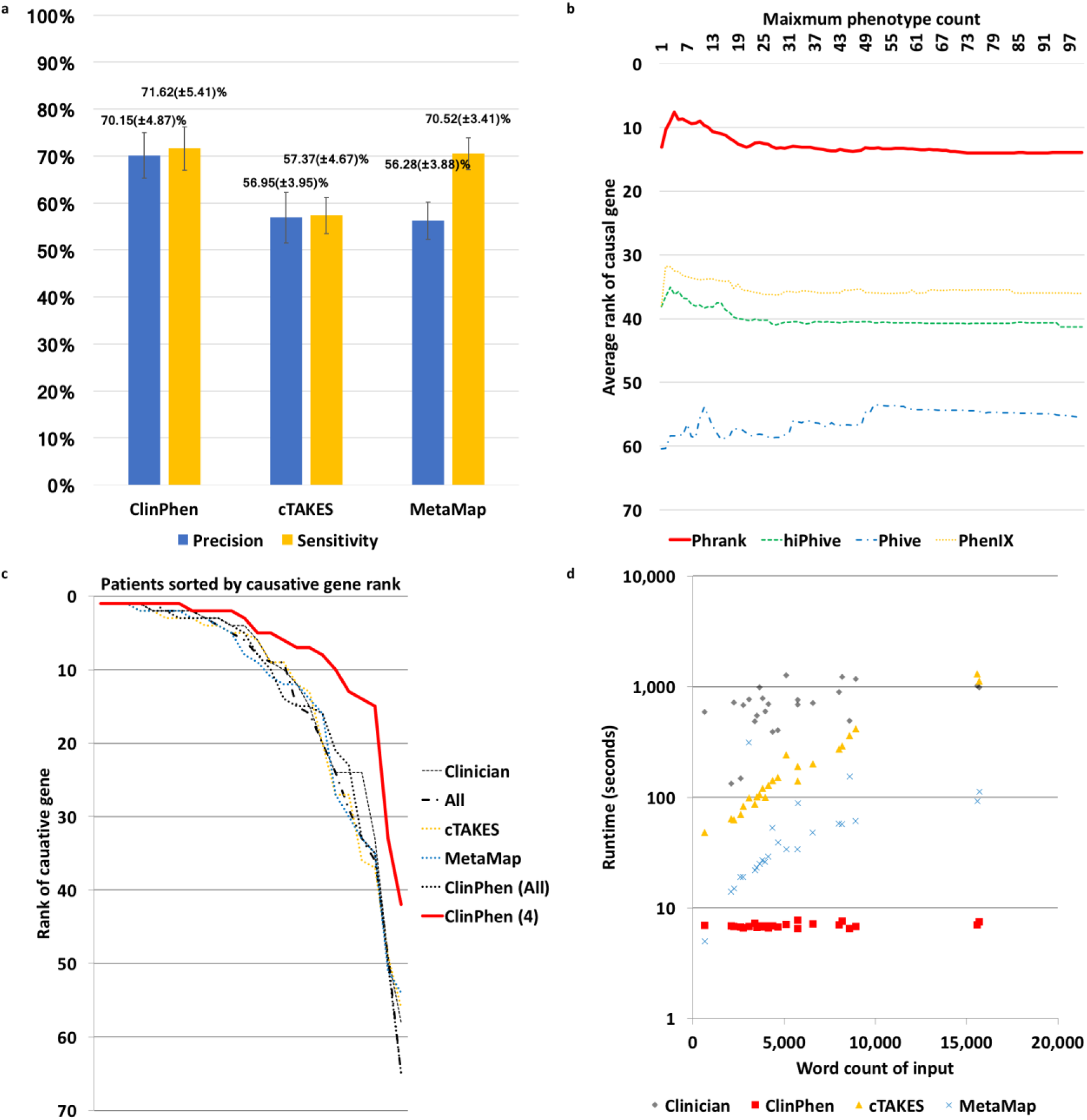
Performance of all extraction methods. (a) Comparison of the extractors’ precision and phenotype sensitivity (higher bars mean higher accuracy). We compared the average precision and sensitivity values of ClinPhen, cTAKES, and MetaMap, using the 24 Test patients as test subjects, and the *All* set (all of the phenotypes found manually and confirmed by a physician to apply to the patient) as the “correct” phenotypes. The average (column) and 95% confidence interval (calculated using Student’s t distribution) of the precision and sensitivity values across the 24 patients are displayed for each extractor. ClinPhen has the highest average precision and sensitivity among the automatic extractors. (b) Causative-gene-ranking performance of each gene-ranking tool when run with different numbers of phenotypes returned by ClinPhen (lower number means better causative gene rankings). ClinPhen was run on the clinical notes of the Test patient set, and the gene-ranking tools were called with the patient’s genetic information and the *n* highest-priority (most-mentioned, first-occurring) extracted phenotypes, with *n* running from 1 to 100 inclusive. The average causative gene rank across the 24 patients was taken for each phenotype-count-limit(*n*)/gene-ranking-tool pairing. The better-performing gene-ranking algorithms rank the causative gene higher when run with a few (around 4) high-priority phenotypes than with all extracted phenotypes. (c) Phrank’s causative-gene-ranking performance across all extraction methods (lower numbers mean better causative gene rankings). We compared the causative gene ranks obtained by running Phrank on the Test patients, with various extracted sets of phenotypes (All manually found, physician-verified phenotypes (*All*) vs. phenotypes considered by a physician to be useful for diagnosis (*Clinician*) vs. automatically extracted phenotypes using various methods). Phrank ranks are sorted lowest-to-highest for each extractor. Phrank performs significantly better when run with ClinPhen’s 4 highest-priority phenotypes (the most-mentioned, earliest-occurring phenotypes in a patient’s clinical notes) than when run with other phenotype sets, manually or automatically extracted. (d) Extractor runtime comparison on each patient (lower number means faster runtime). We measured the runtime of each extractor (ClinPhen, cTAKES, and MetaMap) on each patient’s clinical notes, in seconds. For each patient, we also measured the time taken for a physician to manually scan through the same notes read by the automatic extractors, and encode the phenotypes considered useful for diagnosis to produce the *Clinician* Phenotype set. Each data point is one patient whose clinical notes were scanned by one of the extractors. The horizontal position is the total number of words in the patient’s clinical notes. The vertical position is the time taken for the extractor to run on the notes (logarithmically scaled). While MetaMap’s runtime scales linearly and cTAKES’ exponentially with the total length of the clinical notes, ClinPhen runs in near-constant time. All automatic extraction tools are much faster than manual extraction.

### Automatic extraction of phenotypes accelerates Mendelian disease diagnosis

#### Limiting the number of extracted phenotypes leads to better results with automatic gene-ranking methods

A patient undergoing genome sequencing can have well over 100 candidate genes containing variants of uncertain significance, and each gene takes, on average, an hour to evaluate^2^. Gene-ranking tools expedite the process of finding the causative gene by sorting the genes based on how well their associated symptoms match the patient’s. The closer the causative gene is to rank 1, the sooner clinicians will find it. The rankings are dependent on a provided list of patient phenotypes, meaning that the ideal phenotype set for diagnosis is the one that helps gene-ranking tools rank the causative gene close to the top. We show that this goal is better accomplished not by the full set of patient phenotypes, but by a subset thereof.

For genetic disease diagnosis, a good phenotype set accurately reflects the patient’s symptoms, but a great phenotype set excludes the environmentally caused symptoms, and reflects only the genetically caused ones. Symptoms caused by a common cold, rather than a genetic variant, can easily mislead a gene-ranking tool and make the causative gene harder to identify. ClinPhen, as far as we are aware, is the first phenotype extractor to account for this caveat: phenotypes extracted from the clinical notes are prioritized, first by number of occurrences in the notes (phenotypes that likely pertain to a genetic disease are usually mentioned in multiple clinical notes, and even multiple times in the same note), then by earliest occurrence in the notes (clinicians usually begin a note with a summary of the phenotypes that seem striking and indicative of a genetic disease).

To determine the ideal number of top-priority phenotypes to use when running gene-ranking tools, we ran ClinPhen on the Test patients’ clinical notes, and filtered the extracted phenotypes down to the *n* highest-prioritized phenotypes, for every number *n* from 1 to 100 inclusive. Each set of *n* highest-priority phenotypes was used as input to four automatic gene-ranking algorithms: Phrank^4^, hiPhive^5^, Phive^6^, and Phenix^7^ For each phenotype-count(*n*)/gene-ranking-tool pairing, we found the average causative gene rank across the 24 Test patients (Figure 3b).

The higher-performing gene-ranking tools (Phrank, hiPhive, and PhenIX) generally ranked the causative genes higher at phenotype maxima below 10 (*n* < 10). Phrank, the highest-performing of these, showed a clear spike in performance, yielding the best causative gene rankings at a phenotype maximum of 4 (*n* = 4). It was thus approximated that the 4 highest-priority phenotypes returned by ClinPhen generally lead to the best causative gene rankings.

Across the 24 Test patients, Phrank ranked the causative gene at an average rank of 14.0 with unfiltered ClinPhen phenotypes, and 7.63 with ClinPhen’s 4 top-priority phenotypes (lower number means better ranking). Limiting to the 4 highest-prioritized phenotypes significantly improves Phrank’s causative gene rankings (one-sided Wilcoxon Signed Rank test: p<0.00662).

#### Gene-ranking tools perform better when using automatically extracted phenotypes

Phrank, when run with the 4 phenotypes found and prioritized highest by ClinPhen, puts the causative genes at an average rank of 7.63. If using manually extracted phenotypes were to lead to higher-ranked causative genes, then using ClinPhen-extracted phenotypes would not save time in identifying the causative gene.

We thus set out to show that Phrank does not rank causative genes lower when using ClinPhen’s extracted phenotypes than when using manually extracted phenotypes. We compare two ways of manually extracting phenotypes from clinical notes: manually subsetting all mentioned phenotypes to those that a clinician thinks are most likely to help with the diagnosis (represented by the *Clinician* phenotype set); and listing all mentioned patient phenotypes, whether or not they are likely to help with the diagnosis (represented by the *All* phenotype set). The *Clinician* and *All* phenotype sets were generated for each of the 24 Test patients.

The 24 Test patients were each run through the automatic gene-ranking tool Phrank using each of 6 phenotype sets: the *All* set; the *Clinician* set; ClinPhen(All), the full set of phenotypes returned by ClinPhen; ClinPhen(4), the 4 top-prioritized phenotypes returned by ClinPhen; the phenotypes returned by cTAKES, and the phenotypes returned by MetaMap (Figure 3c). Running Phrank with the *All* set results in an average causative gene rank of 14.3, using the *Clinician* set results in an average rank of 12.9, and using ClinPhen’s 4 top-prioritized phenotypes results in an average rank of 7.63 (lower number means better ranking). Phrank ranked the causative genes significantly higher with the *Clinician* set than with the *All* set (one-sided Wilcoxon Signed Rank test: p<0.0380), and, strikingly, significantly higher with ClinPhen(4) than with the *Clinician* set, or with the phenotypes returned by cTAKES or MetaMap (all p-values < 0.00980). Assuming a clinician examines a ranked gene list from top to bottom, spending an average of one hour evaluating the variants in each gene for their potential to have caused the patient’s phenotypes^2^; using the 4 top-prioritized ClinPhen phenotypes (instead of manually extracted phenotypes) as input to an automatic gene-ranking tool can save roughly 5 hours per case in the diagnostic process.

### ClinPhen consistently runs in less than 10 seconds

A good phenotype extractor runs in a short amount of time. More clinical notes take a longer time to read through, and some patients have far more clinical notes than others do. Therefore, automatic extractors that can quickly extract HPO phenotypes from long collections of clinical notes are ideal.

The Test patients had an average of 4 free-text pre-diagnosis clinical notes per patient. The physician who extracted the *Clinician* set (defined above) of HPO phenotypes from the notes measured the time he took to do so for each patient. For the average patient, he identified 25 phenotypes in 11:55 minutes. Running cTAKES took an average of 4:07 minutes per patient, and running MetaMap took an average of 57 seconds per patient.

The time taken to extract a patient’s phenotypes scaled with the combined length of the notes: for the longest collections of notes, it took over 1,000 seconds to run cTAKES or produce the *Clinician* set, and it took over 100 seconds to run MetaMap. ClinPhen, uniquely, maintained a nearly constant runtime of 7 seconds per patient, even when run on the longest collections of clinical notes. (Figure 3d).

## Discussion

Automatic gene-ranking tools expedite genetic disease diagnosis, but they currently require manually encoding patient phenotypes found into phenotype ontology terms. We show here that an automatic phenotype extractor, ClinPhen, produces an accurate phenotype list in under 10 seconds, and saves an average of 5 hours of candidate gene evaluation per case.

Most of the diagnosis time saved by ClinPhen can be attributed to its unique ability to prioritize the more-relevant extracted phenotypes. Phenotypes that are likely not caused by a genetic disease can derail a diagnosis, and while clinicians use their intuition to filter out these phenotypes, automatic phenotype extractors until now have not done such filtering. When we use a phenotype filter to limit ClinPhen’s phenotypes to the 4 most-mentioned, then earliest-mentioned; automatic gene-ranking algorithms rank the causative gene much higher than they would with unfiltered phenotypes, or even with phenotypes found and filtered by a clinician. ClinPhen thus enables clinicians to search through 8 genes rather than 13 before they find the causative gene, thus reaching a diagnosis an average of 5 hours sooner^2^.

The time saved in phenotype extraction is not to be disregarded, either: while manual extraction, on average, takes 12 minutes of full-time work, clinicians do not tend to extract phenotypes full-time. Since it is a monotonous process, a true manual extraction will involve breaks, and multiple extractions will often be spread out across multiple days. Some patients will be phenotyped this week, some next week, and some a month from now. With ClinPhen, though, a large number of cases can be run through a gene-ranking tool right away, allowing the diagnostic rate to catch up to the accelerating rate and volume of sequence data generation.

Compared to other phenotype extractors, ClinPhen produces more-accurate HPO phenotypes in a shorter amount of time. We optimized ClinPhen for HPO term extraction because HPO terms are commonly used to describe patients with Mendelian diseases^25,38,39^. Rapidly growing databases like OMIM^40^ use HPO terms to describe tens of thousands of disease-phenotype associations. ClinPhen could be used to accelerate the growth of these databases by quickly analyzing patients’ clinical notes and finding new disease-phenotype associations at a rate unachievable by clinical experts, or even by other phenotype extractors.

Upon publication, ClinPhen will be made available at bejerano.stanford.edu/clinphen, as a standalone, free-to-use, noncommercial tool. Users can download ClinPhen and run it on a patient’s clinical notes to get an encoded list of HPO terms, along with the number of mentions and location of the earliest mention of each phenotype. The large number of undiagnosed patients^11^ with presumed Mendelian diseases necessitates a rapid diagnostic process. With the help of ClinPhen, clinicians can accurately diagnose patients more than 5 hours ahead of schedule, and provide them with faster, more-effective medical care.

## Acknowledgments

We thank Stanford Children’s Health for providing the medical records, and Julia Buckingham for assistance with obtaining the patient data. We also thank members of the Bejerano Lab and Elijah Kravets for their feedback on the project. CD was supported by a Bio-X Undergraduate Summer Research Program (run by Heideh Fattaey). JB was supported by a BioX SIGF fellowship. GB was supported by DARPA and the Stanford Pediatrics Department.

## Author contributions

CD wrote the extraction program under the supervision of JB and GB. CD, JB, JAB, and GB all provided ideas for improving and testing the extractor. CD and JB identified undiagnosed and diagnosed patient cases. JAB confirmed the diagnoses. CD downloaded patient records. CD produced the *All* phenotype sets, and EEB verified them. EEB produced the *Clinician* phenotype sets. CD ran tests (designed by CD, JB, and GB) on the extractor for precision, sensitivity, and automatic gene ranking performance. CD, JB, and GB wrote the manuscript. JB, CD and GB designed the study. All authors read and commented on the manuscript.

## Web Resources

Upon publication, ClinPhen will be made publicly available at bejerano.stanford.edu/clinphen as a non-commercial, free-to-download tool.

